# Leveraging Light Detection and Ranging (LiDAR) to Elucidate Forest Structural Conditions that Influence Eastern Whip-poor-will Abundance

**DOI:** 10.1101/2025.06.26.661842

**Authors:** Jeffery T. Larkin, Malcolm Itter, Cameron J. Fiss, Lauren M. Chronister, Justin Kitzes, Jeffery L. Larkin, Halie Parker Larkin, Darin J. McNeil, Anthony W. D’Amato, Michael E. Akresh, David I. King

## Abstract

Eastern North American forests are degraded due to land use history and are threatened by numerous factors that further reduce their structural complexity, which contributes to population declines of many taxa. As such, many agencies and their conservation partners are employing habitat centric conservation efforts. Increased availability of airborne Light Detection and Ranging (LiDAR) data provides an opportunity to quantify fine-scale structural habitat characteristics for forest wildlife. One such species of conservation concern, the eastern whip-poor-will (*Antrostomus vociferus*), requires diverse forest structural conditions to meet its breeding season habitat requirements. We used airborne LiDAR data and autonomous recording units (ARUs) to identify elements of forest structure that influence whip-poor-will breeding season abundance in Pennsylvania, USA. Specifically, we applied a machine-learning classifier for whip-poor-will song to audio recordings obtained from 851 ARUs that were deployed in forested landscapes and then created daily detection histories to estimate whip-poor-will relative abundance. Whip-poor-wills were detected at 334 survey locations (41%). Abundance exhibited positive linear relationships with percent forest cover and percent oak forest and a negative linear relationship with percent impervious cover. Whip-poor-will abundance was also influenced by forest structure, with abundance exhibiting a quadratic relationship with two LiDAR-derived covariates; canopy heterogeneity and height within 300 m. Using these results, we predicted whip-poor-will abundance and habitat management potential. Whip-poor-will conservation in our study region will depend on public and private land efforts that maintain heavily forested, oak dominated landscapes that are managed using practices that increase canopy height diversity among and within stands.

## Introduction

Forests of eastern North America (hereafter, eastern forests) host among the highest levels of biodiversity in the Nearctic realm (Marshall, 2006; Petrides, 1972; Sibley, 2014). This diversity is, in part, the result of millennia of forest dynamics that generated structural complexity across the landscape (Ellsworth and McComb, 2003; MacCleery, 2011; Oliver and Larson, 1996).

Indeed, the biodiversity of eastern forests coevolved under conditions that resulted from a continuous interplay between ecological succession and disturbances that varied in frequency and severity, such as fires (Abrams, 1992; Nowacki and Abrams, 2008), weather events (MacCleery, 2011), and stand-disturbing wildlife (Greenberg and Collins, 2015, Ellsworth and McComb, 2003). Structurally complex forests are important to many taxa (Divoll et al., 2022; McNitt et al., 2020; Mathis et al., 2021) and are known to promote biodiversity (Phillips, 2011).

Unfortunately, eastern forests are experiencing unprecedented challenges that degrade their conditions and threaten associated biodiversity (Shifley et al., 2014). Most contemporary eastern forests originated following widespread, intensive land uses associated with European colonization (e.g., agricultural clearing, clearcut harvesting) and often have lower compositional and structural complexity than those pre-settlement (Nowacki and Abrams, 2008, MacCleery, 2011). Factors such as the suppression of select disturbances (Hanberry et al., 2020; Nowacki and Abrams, 2008; Sabadosa, 2021), excessive deer browse (Parker et al., 2020), invasive species (Morin and Liebhold, 2016; Ward et al., 2018), and unsustainable logging practices (Curtze et al., 2022) are contributing to the continued degradation of eastern forests. Forest degradation threatens the persistence of certain plant community types (e.g., oaks [*Quercus* spp.]; Dey, 2014; USDA Forest Service, 2024), and may lead to reduced structural complexity within stands and across forest landscapes (Schulte et al., 2007). Thus, species groups such as eastern forest birds, that are known to benefit from heterogenous structural conditions, are experiencing precipitous population declines (Rosenberg et al., 2019; Zilkowski et al., 2024). In turn, most state and federal agencies have made efforts to enhance or create habitat for declining species through active forest management (Bakermans et al., 2011; Wood et al., 2013; Lambert et al., 2017).

Developing effective habitat management guidelines for forest birds is challenging because they select resources at a variety of spatial scales, from landscape-level territory placement decisions to micro-site determinations such as where to forage, roost, and nest (Chandler et al., 2012; Fiss et al., 2020; Fuoco et al., 2024). Advances in wildlife monitoring techniques (e.g., autonomous recording units [ARUs]) have allowed conservation scientists to better assess factors that influence species’ occurrence and demography across large spatial extents (Chronister et al., 2024; Larkin et al., 2024b). Yet, identifying associations with fine-scale vegetation structure remains challenging due to limitations in collecting such data, especially across a large quantity of survey locations.

Light Detection and Ranging (LiDAR) derived datasets have the potential to help researchers better understand associations between wildlife and fine-scale forest structure at multiple spatial extents ranging from within-stand to regional landscapes (Goetz et al., 2010; Bulluck et al., 2022; McNeil et al., 2023). Recent research using LiDAR derived metrics has found important avian associations with structural conditions at spatial extents beyond what traditional field-based vegetation surveys capture (McNeil et al., 2023; Larkin et al., 2024a). Furthermore, studies have demonstrated that LiDAR derived metrics are more effective at identifying structural associations than field collected measures (McNeil et al., 2023) and other publicly available datasets (e.g., Multi-Resolution Land Characteristics National Land Cover Database Products [Dewitz, 2023]; Bulluck et al., 2022). Given these findings, LiDAR can greatly advance the conservation of many declining wildlife species (e.g., McNeil et al., 2023; Fuoco et al., 2024), especially those influenced by habitat factors beyond the stand scale (e.g., eastern whip-poor-will, *Antrostomus vociferus*; Vala et al., 2020; Larkin et al., 2024b).

The eastern whip-poor-will (hereafter, whip-poor-will) is a migratory nightjar that breeds in eastern North America (Cink et al., 2020). Whip-poor-will populations have been experiencing significant population declines (Ziolkowski et al., 2024, NABCI, 2025) and are listed as species of greatest conservation need on many state wildlife action plans (e.g., Pennsylvania, PGC-PFBC, 2015; Massachusetts, MA-DFW, 2015; Virginia, VA-DGIF, 2015; etc.). Whip-poor-will population declines are thought to be driven, in part, by loss of early successional conditions on the breeding grounds (NABCI, 2025). A considerable body of research has indicated that whip-poor-will occupancy and abundance are influenced by site and landscape factors during the breeding season (e.g., basal area and percent impervious cover, respectively; Spiller and King, 2021; Souza-Cole et al., 2022; Larkin et al., 2024b). Indeed, past research has been important for informing whip-poor-will breeding season conservation and management efforts. However, an understanding of how forest structure effects whip-poor-will abundance at large spatial scales is currently lacking from the literature.

Here, we conducted an analysis of whip-poor-will associations with forest structure informed by recently collected airborne LiDAR data across the state of Pennsylvania. To our knowledge, this is the first study that leverages LiDAR to better understand habitat associations for this species. More broadly, this research demonstrates the value of incorporating LiDAR data in analyses to identify wildlife-habitat relationships that would otherwise not be feasible to examine. Our study objectives were to: 1) identify forest structural characteristics associated with high whip-poor-will abundance; and 2) create a fine-resolution, predictive map of whip-poor-will abundance in Pennsylvania to identify critical areas for conservation and future habitat management efforts. We hypothesized that whip-poor-will abundance would be greatest at locations with high levels of horizontal complexity, specifically forested landscapes with diverse successional conditions, due to the species’ use of forest edges (Cink et al., 2020; Wilson and Watts, 2008; Grahame et al., 2021). Further, we postulated that locations with an interspersion of mature and intermediate amounts of early successional forests would have the highest abundances given the species need for both young and mature forests (Akresh et al., 2016; Larkin et al., 2024b). Given that forest structure is only one of many factors that could influence whip-poor-will abundance, we also considered several landscape variables in our analysis. We hypothesized abundance would have a positive relationship with forest cover (Souza-Cole et al., 2022; Thompson et al., 2022) and oak dominated forest types (Larkin et al., 2024b), and a negative relationship with impervious cover (Souza-Cole et al., 2022; Larkin et al., 2024b).

## Methods

### Study Area

We studied whip-poor-will abundance in public and private forests across Pennsylvania, USA (Fig. 1). Forest stands included in our study represented a diversity of structural conditions ranging from recent clearcuts and partial timber harvests to mature, closed-canopy forests. Most stands were classified as either oak-hickory (e.g., *Carya* spp., *Quercus* spp., *Pinus* spp.) or northern hardwoods (e.g., *Acer* spp., *Betula* spp., *Fagus grandifolia*, *Tsuga canadensis*).

**Figure 1.**
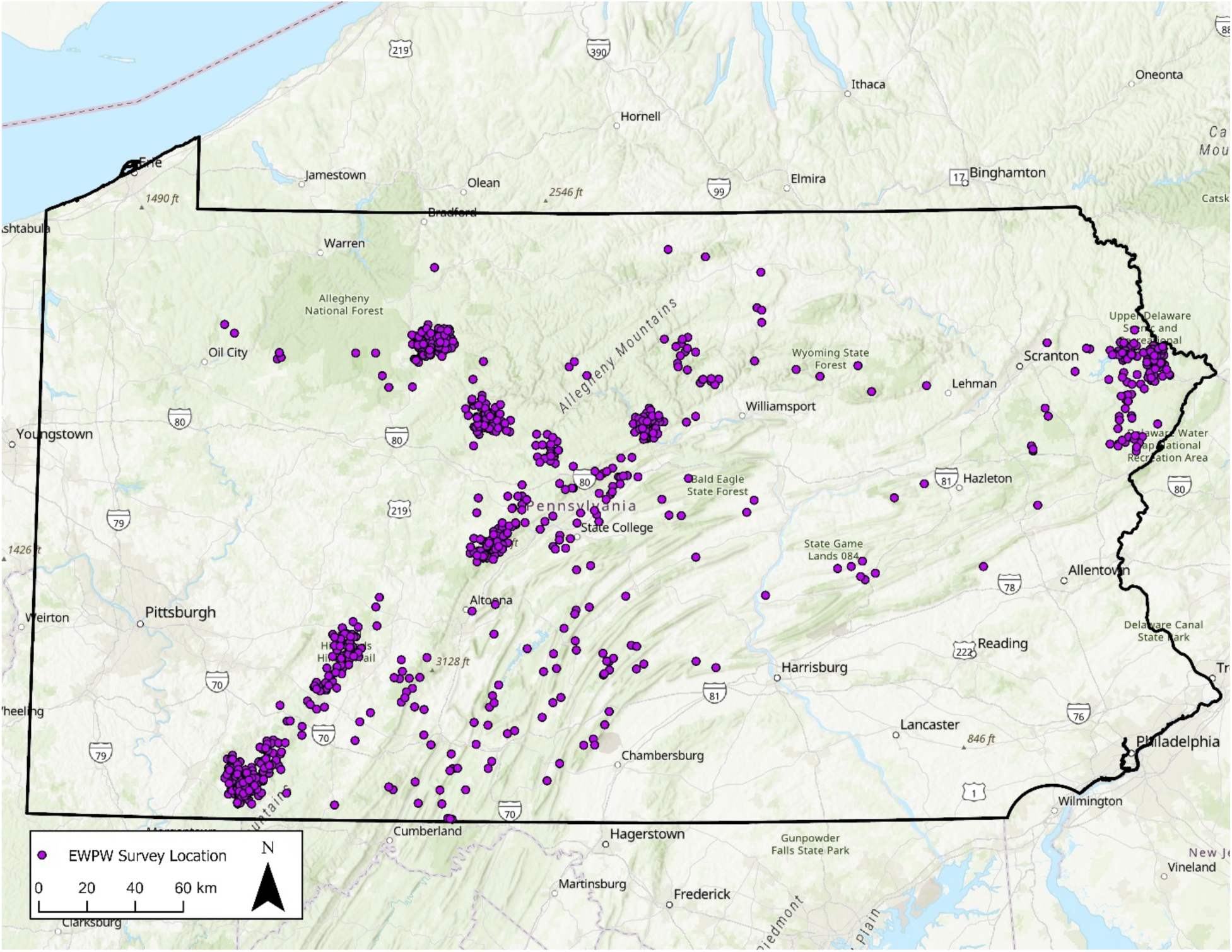
Map displaying the 851 survey locations (purple points) where autonomous recording units (ARUs) were deployed across Pennsylvania, USA in 2020 and 2021. ARUs were deployed to monitor actively and passively managed sites for territorial eastern whip-poor-will (*Antrostomus vociferus*). Data from these units were used in subsequent abundance analysis which aimed to identify remotely sensed landscape and forest structural variables that influenced eastern whip-poor-will abundance. Note: Each private land survey location was shifted in a random direction 0–25 km to preserve landowner privacy.

Depending on disturbance history, understories ranged from sparsely vegetated, to young regenerating forests characterized by a mix of herbaceous (e.g., *Solidago* spp., graminoids, and ferns) and woody vegetation (e.g., shrubs, saplings, and brambles) with varying heights and densities. Elevation ranged between 197 and 884 m above sea level. For additional information on study area and forest management treatments, see Larkin et al. (2024b).

### Whip-poor-will Survey Locations

To establish whip-poor-will survey locations within pre-selected stands for monitoring, we used the “Create Random Points” tool in ArcGIS Pro Version 2.9.1 (ESRI, 2021). This placed random survey locations, stratified by our pre-selected monitoring stands representing a gradient of management intensity. We ensured that all survey locations were ≥50 m from the treatment edges (to limit edge effects) and spaced a minimum of 500 m apart to maintain spatial independence (Bibby et al., 2000; Larkin et al., 2024b). Applying the above criteria, we generated 851 unique survey locations across 152 properties (23 public land units and 129 private properties; Fig. 1). This included 211 locations on private forests enrolled in Natural Resource Conservation Service’s (NRCS) conservation programs [NRCS’s Regional Conservation Partnership Program (RCPP) for cerulean warbler (*Setophaga cerulea*, CERW; n = 69) and Working Lands for Wildlife (WLFW) program for golden-winged warbler (*Vermivora chrysoptera*, GWWA; n = 142)], and 640 locations on public forests managed by the Pennsylvania Game Commission (PGC), Department of Conservation and Natural Resources State Parks (DCNR-State Parks) and State Forests (DCNR-State Forests).

### Autonomous Recording Units and Acoustic Data Processing

We used ARUs (AudioMoths, Open Acoustic Devices) to collect audio recordings at each survey location from late April-July in 2020 or 2021 (Hill et al., 2019; Larkin et al. 2024b). ARUs were attached to a woody stem 1.5-2 m from the ground and programmed to record for 2 hours after sunset (2100-2300 EST). Collected audio recordings were processed using OpenSoundscape version 0.6.1 in Python (Van Rossum and Drake, 1995; Lapp et al., 2023). Recordings were broken into clips and then processed through a binary, single target automated classifier. The classifier assessed each clip for the presence of the whip-poor-will song and assigned a score, which was indicative of the classifier’s confidence that a given clip contained whip-poor-will song. Using a threshold approach (precision = 1.00; recall = 0.35 at a threshold of 4.30), we created a daily detection history for each survey location whereby “1” denoted a whip-poor-will detection and “0” denoted a non-detection. For additional information on ARUs and acoustic data processing, see Larkin et al. (2024b).

### LiDAR

To quantify aspects of forest structure, we used LiDAR metrics (10 m resolution) developed by Fisher et al. (2024), which were derived from Pennsylvania’s publicly available LiDAR point clouds (https://apps.nationalmap.gov/lidar-explorer/). These LiDAR data were collected via aircraft over multiple campaigns spanning spring 2017 through spring 2020 prior to forest leaf-out. The LiDAR-derived metrics included in our analysis were those that we expected to be predictive of whip-poor-will territory placement, and related to forest structural characteristics that can be influenced by management activities. Specifically, we used the following LiDAR metrics: 1) height below which 95% of returns occurred (“p95”; a measure of canopy height); and 2) standard deviation of p95 (hereafter, “SD p95”; a measure of canopy height variability). We summarized LiDAR values within 300 m radii (30 ha; mean size of a whip-poor-will home range in Pennsylvania; Notarianni, *unpublished data*) of each survey location using the *Focal Statistics* tool in ArcGIS Pro (ESRI, 2021). Specifically, we used the argument “circle” for the neighborhood with a radius of 30 and unit type of “cell” (10 m cell x 30 cells = 300 m radius focal raster). We used statistics type “Mean” and “Standard Deviation” to create the p95 and SD p95 focal rasters, respectively. LiDAR acquisition for a portion of northwest Pennsylvania was flawed and resulted in low quality data (see Fisher et al., 2024). Additionally, LiDAR was not acquired for two counties (York and Chester) in southeast Pennsylvania. Thus, we removed these portions of Pennsylvania from our analysis. Further, we dropped any survey locations that were within, or less than 300 m from the edge of, an area without LiDAR (n = 5). Using the *extract()* function from the package “terra” in program R (R Core Team, 2021; Hijmans, 2023), we extracted the value at each survey location (Table 1).

**Table 1.** Summary statistics for all variables considered for inclusion in Royle-Nichols models to estimate eastern whip-poor-will (*Antrostomus vociferus*) abundance in Pennsylvania, USA between 2020 and 2021.

| Covariate | Type | Description | Units | Min | Max | Mean | Median |
| --- | --- | --- | --- | --- | --- | --- | --- |
| MR | Detection | Moon above the horizon (1/0) | NA | 0 | 1 | NA | NA |
| Day |  | Day of survey window | days | 1 | 15 | NA | NA |
| p95 | State | Canopy height | meters | 3.93 | 31.86 | 19.34 | 19.34 |
| SD p95 |  | Variation in canopy height | meters | 1.24 | 13.00 | 5.40 | 5.11 |
| Oak |  | Precent oak dominated forest | percent | 0.00 | 100.00 | 78.72 | 93.97 |
| Forest |  | Precent forest cover | percent | 45.74 | 100.00 | 95.84 | 98.91 |
| Impervious |  | Precent impervious cover | percent | 0.00 | 11.00 | 0.25 | 0.03 |

### Landscape Analysis

Given that LiDAR-derived forest structural metrics are not the only factors needed for predicting and mapping whip-poor-will abundance, we included several other remotely-sensed variables that describe landscape characteristics at scales known to influence whip-poor-wills.

Specifically, we included the percent total forest cover within 1500 m (Vala et al., 2020; Thompson et al., 2022); percent oak forest (oak-hickory and oak-pine) within 1500 m (Larkin et al., 2024b); and mean percent impervious within 500 m (Larkin et al., 2024b).

Using program R (R Core Team, 2021) and the 2019 US Forest Service forest type groups dataset (30 m resolution; USDA Forest Service, 2023), we created a binary raster whereby oak-hickory and oak-pine forest types were designated as “1” and all other forest types were “0”. Using this binary oak/non-oak raster and the *Focal Statistics* tool in ArcGIS Pro (ESRI, 2021), we created a new raster (30 m resolution) that used a moving window to calculate the percent oak forest within 1500 m. We used this same process to create the percent forest raster with the Dynamic World dataset from Google Earth Engine (Brown et al., 2022). We classified “Trees” (class 1) and “Shrub & Scrub” (class 5) as forest, “1”, and all other classes as non-forest, “0”, to create a binary forest raster (10 m resolution), which was used to calculate focal statistics at 1500 m. The process for calculating mean percent impervious was similar, however the creation of a binary raster was not necessary since the impervious dataset (Dewitz, 2023) is continuous rather than categorical. Using these rasters (percent oak and forest within 1500 m and mean percent impervious within 500 m) and the *extract()* function in the package “terra” (R Core Team, 2021; Hijmans, 2023), we extracted the value at each survey location (Table 1).

### Data Analysis

We applied a Royle-Nichols (R-N) model (Royle and Nichols, 2003) to assess the factors affecting whip-poor-will abundance fit using the “unmarked” package in R (Fiske and Chandler, 2011; Kellner et al., 2023). R-N models have previously been shown to provide reliable estimates of avian abundance using machine-learning generated detection histories when validated to ensure no false positives (Fiss et al., 2024). In the context of ARU collected data, where the true listening radius of the recorder is unknown, it is best to consider estimates from R-N models as a relative index of abundance. Nevertheless, these models allowed us to achieve our primary interest in comparing abundance among survey locations and assessing impact of forest structure on abundance.

Daily presence/absence of whip-poor-will for a 10-day window, which fell in the Nightjar Survey Network’s 2020 and 2021 survey dates (nightjars.org), was used to generate a detection history at each sampling location. This detection history served as the response variable in the R-N model. As such, any survey locations with <10 days of recording were excluded from analyses (n=35). To account for imperfect detection we considered two detection covariates in our analysis: 1) Moon above the horizon (“MR”); and 2) Day of the survey window (“Day”; Table 1). MR, a binary variable, was calculated using the website *Time and Date* (timeanddate.com), which provides information about the lunar cycle. Day was calculated using the Nightjar Survey Network’s survey dates (nightjars.org), whereby day one corresponded with the first survey date of a given window and continued until the end of the window. These are two variables known to influence whip-poor-will detection (Wilson and Watts, 2006). We did not explicitly model lunar phase or length of time the moon was above the horizon for each recording because both have a relationship with day of the survey window. Rather, we modeled both linear and quadratic effects of Day because lunar phase peaked in the middle of the survey window and length of time the moon was above the horizon decreased linearly.

Before building models, we tested for correlation among detection and state covariates by calculating pairwise Pearson’s Correlation coefficients. Neither of the detection covariates, nor the five state covariates were correlated (correlation coefficient <±0.5 for all covariates considered; Sokal and Rohlf, 1969). To identify the detection covariates that would be carried into our model sets, we tested four detection-only models applying a forward selection approach (Table 2). We included quadratic terms for both LiDAR metrics, because published literature suggests that these structural metrics best explain whip-poor-will distributions at intermediate values (Wilson and Watts, 2008; Spiller and King, 2021). Once detection covariates were identified, we tested six candidate models applying a forward selection approach (Burnham and Anderson, 1998). Covariates were added in order of importance to whip-poor-will ecology based on past literature and our knowledge of the species. A covariate was retained in subsequent models only if its inclusion resulted in a lower AIC_c_. We considered covariates for which coefficient 95% confidence intervals did not overlap with zero to be informative predictors (Chandler et al. 2009). To assess model fit, we ran a MacKenzie and Bailey goodness-of-fit test (*mb.gof.test()*) with 1000 simulations (Kéry and Royle, 2015) applied to our top model.

**Table 2.** Four detection-only and five full models tested by applying a forward selection approach in an analysis that used remotely sensed data (autonomous recording units, LiDAR, and landcover data) to estimate eastern whip-poor-will (*Antrostomus vociferus*) abundance across Pennsylvania, USA between 2020 and 2021. Variables considered in the detection-only model were: MR (1/0; moon above the horizon) and Day (day of the survey window). State variables considered in the full models were: average p95 (canopy height) and SD p95 (variation in canopy height) within 300 m, percent oak forest and percent forest cover within 1500 m, and percent impervious cover within 500 m of survey locations.

| | Model Name | k | AICc | ΔAICc | CumWt. | LL |
| --- | --- | --- | --- | --- | --- | --- |
| Detection-only | $\Psi(\sim \text{MR} + \text{Day} \sim 1)$ | 4 | 4585.30 | 0.00 | 1.00 | -2288.6 |
| | $\Psi(\sim \text{MR} \sim 1)$ | 3 | 4601.45 | 16.15 | 1.00 | -2297.7 |
| | $\Psi(\sim \text{Day} + \text{Day}^2 \sim 1)$ | 4 | 4762.37 | 177.07 | 1.00 | -2377.2 |
| | $\Psi(\sim \text{Day} \sim 1)$ | 3 | 4854.83 | 269.54 | 1.00 | -2424.4 |
| | $\Psi(\sim 1 \sim 1)$ | 2 | 5093.55 | 508.25 | 1.00 | -2544.8 |
| Full Models | $\Psi(\sim \text{MR} + \text{Day} \sim \text{p95} + \text{p95}^2 + \text{SD p95} + \text{SD p95}^2 + \text{Oak} + \text{Impervious} + \text{Forest})$ | 11 | 3989.53 | 0.00 | 0.91 | -1983.6 |
| | $\Psi(\sim \text{MR} + \text{Day} \sim \text{p95} + \text{p95}^2 + \text{SD p95} + \text{SD p95}^2 + \text{Oak} + \text{Impervious})$ | 10 | 3994.11 | 4.57 | 1.00 | -1986.9 |
| | $\Psi(\sim \text{MR} + \text{Day} \sim \text{p95} + \text{p95}^2 + \text{SD p95} + \text{SD p95}^2 + \text{Oak})$ | 9 | 4056.97 | 67.44 | 1.00 | -2019.4 |
| | $\Psi(\sim \text{MR} + \text{Day} \sim \text{p95} + \text{p95}^2 + \text{p95STD} + \text{SD p95}^2)$ | 8 | 4105.08 | 115.55 | 1.00 | -2044.5 |
| | $\Psi(\sim \text{MR} + \text{Day} \sim \text{p95} + \text{p95}^2)$ | 6 | 4169.52 | 179.99 | 1.00 | -2078.7 |
| | $\Psi(\sim 1 \sim 1)$ | 2 | 5093.55 | 1104 | 1.00 | -2544.8 |

We applied the final model to predict whip-poor-will abundance across Pennsylvania based on conditions at the time of data collection. Additionally, in R, we calculated the difference between predicted abundance under observed conditions and predicted abundance after setting forest structural variables at the values that maximized abundance. The difference between predicted whip-poor-will abundance under observed and ideal forest structural conditions was used as a measure of the potential for forest management, assuming the goal of management is to create or enhance whip-poor-will habitat. Lastly, we used ArcGIS Pro (ESRI, 2021) and conservation land boundary layers (PA GeoData, 2025) to quantify the area classified as low, medium, and high predicted abundance and management potential by forest ownership type (e.g., federal, state, and private). To do this, we reclassified both rasters into three categories. The predicted abundance raster categories were: low (0.10 – 1.00 individuals), medium (>1.00 – 3.00 individuals), and high (>3.00 individuals). The management potential raster categories were: low (change in abundance <2.00), medium (change in abundance 2.00 – 3.99), and high (change in abundance ≥4.00).

## Results

Whip-poor-will were detected at 334 out of 811 survey locations (mean number of detections over ten-day period = 2.9, SD = 3.8). Our top ranked model contained both detection covariates (MR and Day), linear and quadratic terms for both LiDAR covariates (p95 and SD p95), and three landscape covariates (percent oak, percent forest, and mean percent impervious; Table 2). The overdispersion parameter (ĉ) was 1.03, indicating no substantial overdispersion. As such, AIC_c_ was used for model comparison. Detection probability (*p*) was highest when the moon was above the horizon (mean *p* = 0.45), compared to when it was below (mean *p* = 0.05; Table 3).

**Table 3.**
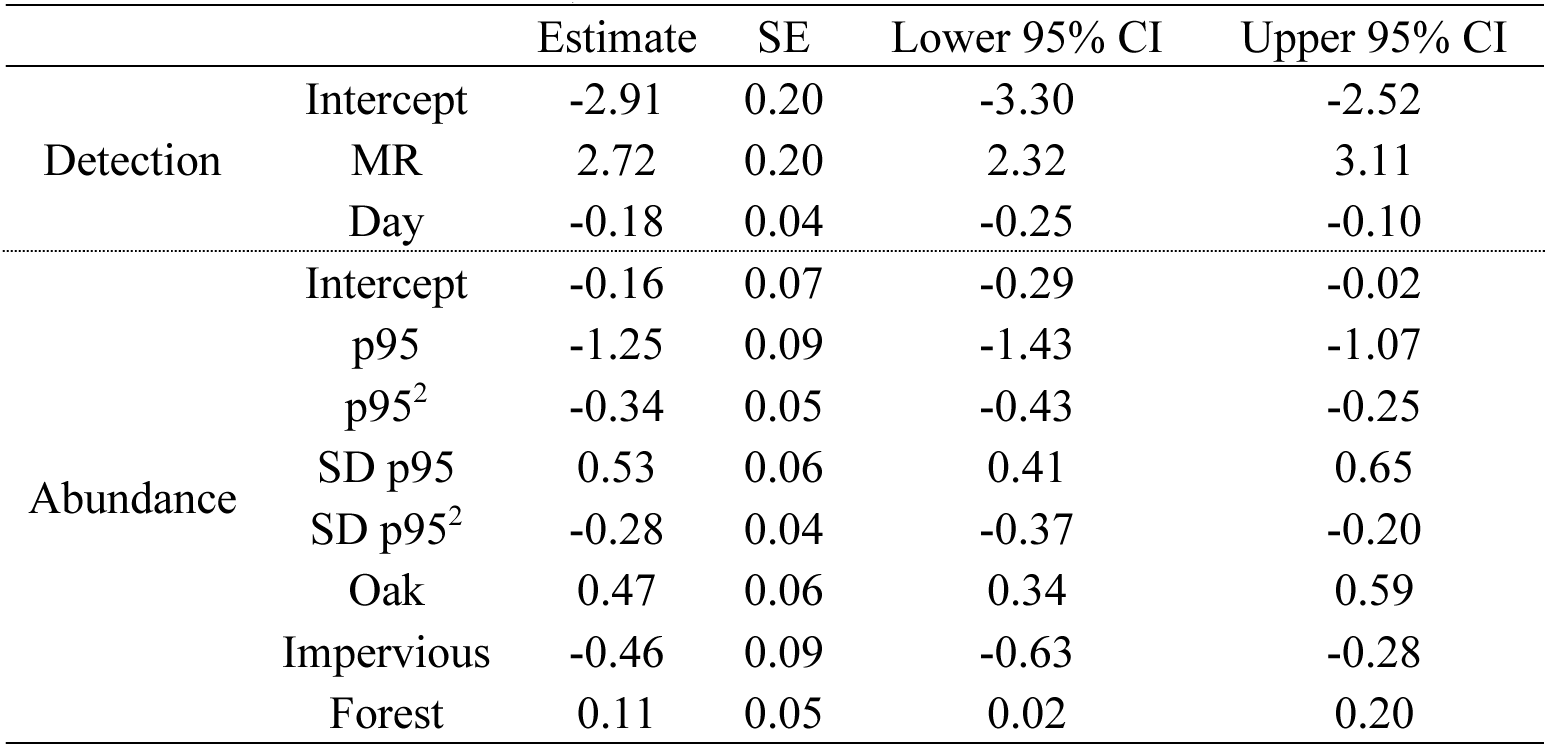
Summary of the top model in an analysis that used remotely sensed data (autonomous recording units, LiDAR, and landcover data) to estimate eastern whip-poor-will (*Antrostomus vociferus*) abundance across Pennsylvania, USA between 2020 and 2021. Detection variables in the model were: MR (1/0; moon above the horizon) and Day (day of the survey window). State variables were: average p95 (canopy height) and SD p95 (variation in canopy height) within 300 m, percent oak forest and percent forest cover within 1500 m, and percent impervious cover within 500 m of survey locations. All variables were considered biologically significant (95% confidence intervals do not include 0).

Similarly, detection had a negative relationship with Day (Table 3). Detection was highest on day one (mean *p* = 0.52) and lowest on day 15 (mean *p* = 0.32) when the moon was above the horizon. When the moon was below the horizon, detection decreased greatly across all days but held the same negative linear relationship (detection probability 0.07 on day one, and 0.03 on day 15).

There was a concave relationship between mean whip-poor-will abundance and p95 (canopy height) and SD p95 (canopy height variability). Specifically, mean abundance increased as a function of p95 and was maximized at 10.1 m (Fig. 2A; Table 3). Similarly, mean abundance increased as a function of SD p95 and was maximized at 7.4 m (Fig. 2B; Table 3).

**Figure 2.**
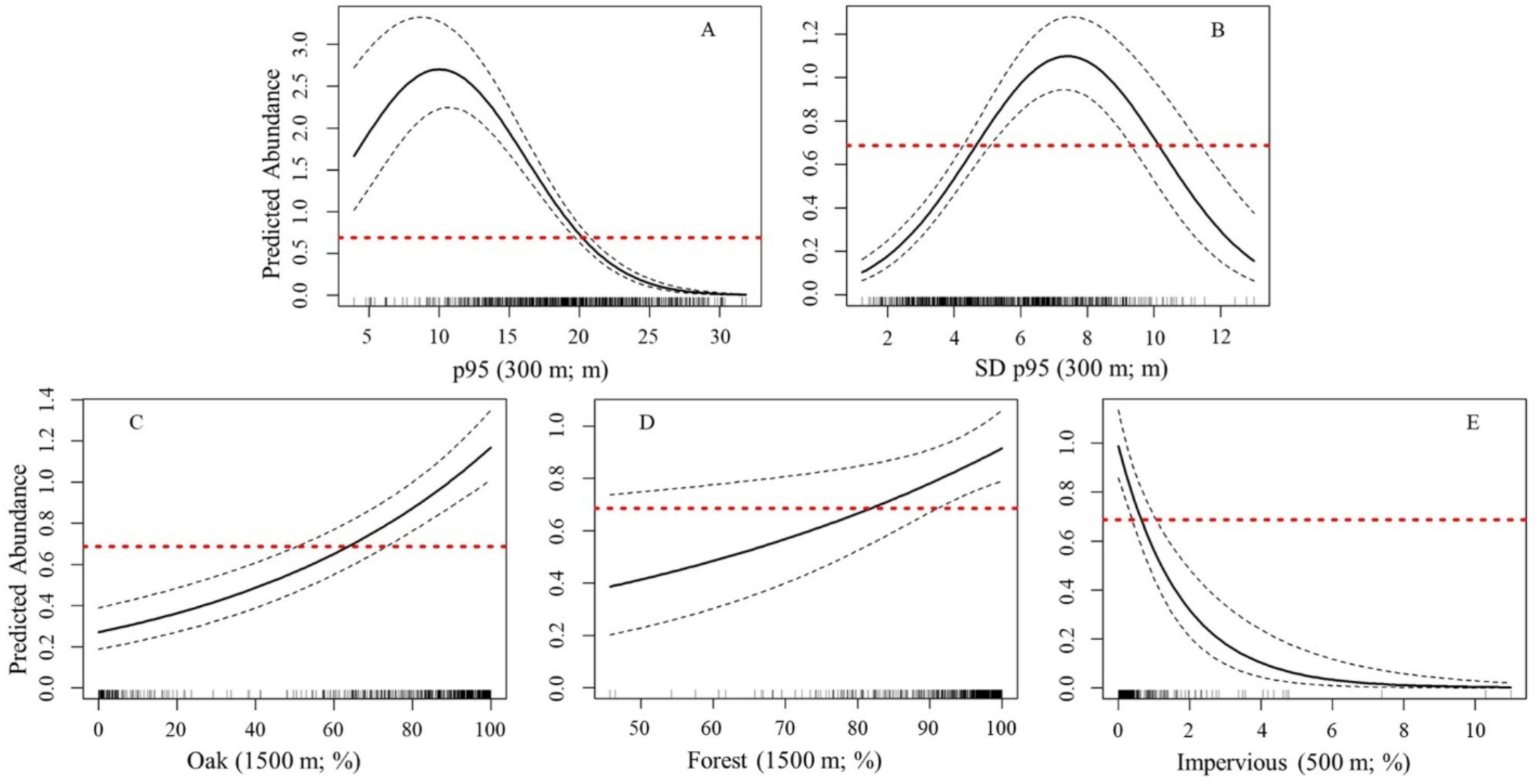
Predicted abundance of eastern whip-poor-will (*Antrostomus vociferus*) in relation to two LiDAR-derived (p95 [canopy height] and SD p95 [variation in canopy height]), and three landscape (percent forest, oak, and impervious), covariates included in a study conducted in Pennsylvania, USA between 2020 and 2021. The solid black line is the model-predicted trendline, the dotted black lines are the upper and lower 95% confidence intervals, and the red dotted line is mean predicted eastern whip-poor-will abundance. X-axis label contains the spatial extent at which the variable best predicted eastern whip-poor-will abundance and the unit in parenthesis, respectively.

Further, there was a significant positive relationship between mean whip-poor-will abundance and both percent oak forest (Fig. 2C; Table 3) and overall forest cover (Fig. 2D; Table 3) within 1500 m and a negative relationship with percent impervious within 500 m (Fig. 2E; Table 3).

LiDAR point clouds of survey locations indicate that high abundance sites contained canopy height diversity within 300 m, while those with few whip-poor-will were homogenous closed canopy forests (Fig. 4).

**Figure 4.**
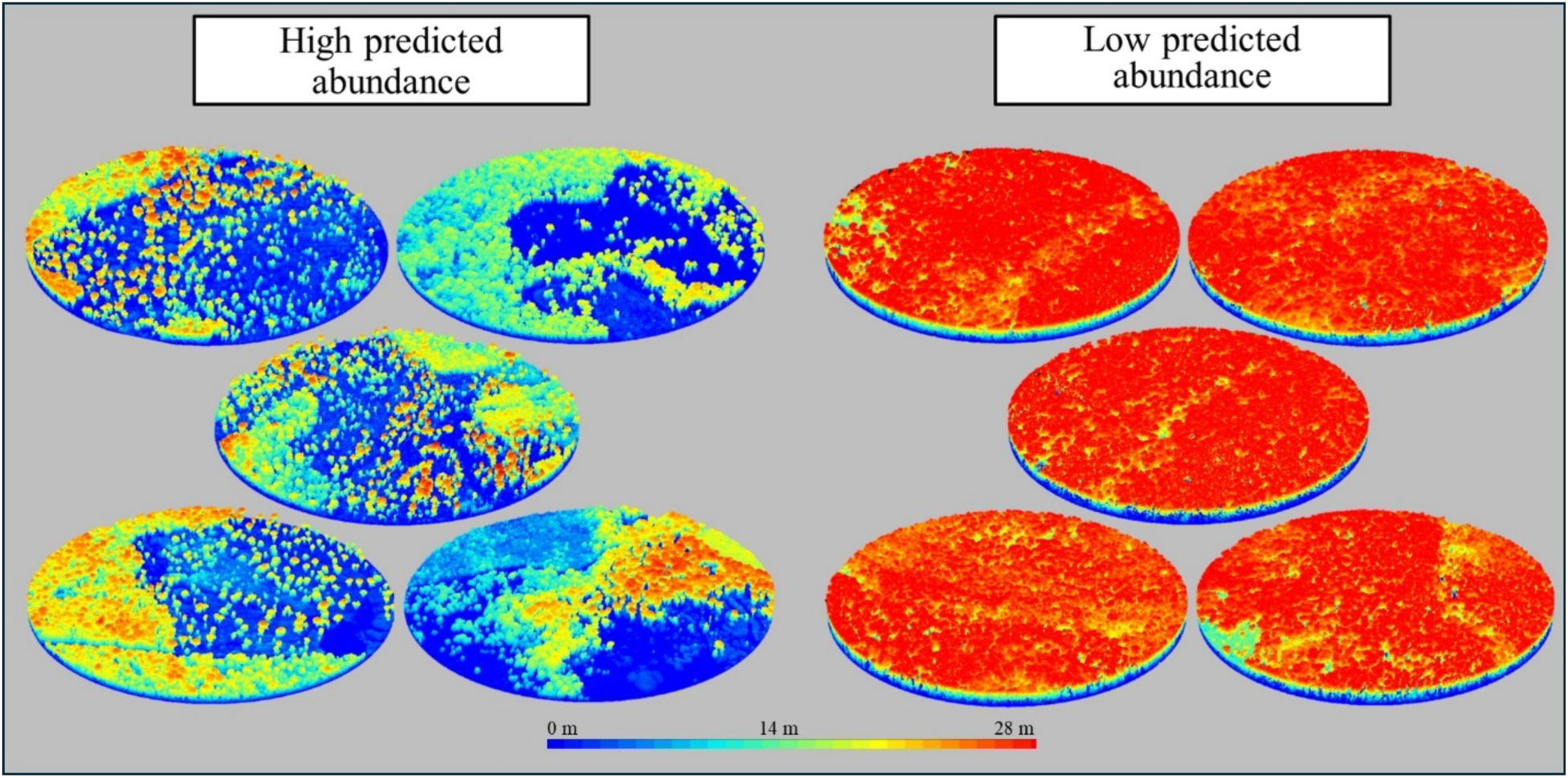
LiDAR cookies, 300 m radius, for the top five highest and lowest predicted eastern whip-poor-will (*Antrostomus vociferus*) abundance survey locations from a study in Pennsylvania, USA between 2020 and 2021. Predicted abundances ranged from 0 – 5.64. Blue and red areas represent short (e.g., saplings and brambles) and tall (e.g., canopy trees) vegetation, respectively.

Predicted whip-poor-will abundance varied considerably throughout our study area with the highest estimates concentrated in the central portion of the state and consistently low estimates across the northern portion (Fig. 3). Our assessment of whip-poor-will predicted abundance relative to ownership type reveals that private forests host approximately 73, 79, and 64 percent of low, medium, and high predicted abundances, respectively (Table 4). Collectively, forests managed by state agencies account for approximately 25, 20, and 36 percent of low, medium, and high predicted abundances, respectively (Table 4). Similar to predicted abundance, the forests of central Pennsylvania have the greatest management potential, while those in northern portions have limited potential (Fig. 5). Private forests account for 84, 67, and 43 percent of areas identified as having low, medium, and high management potential, respectively (Table 4). Collectively, forests managed by state agencies account for approximately 13, 32, and 56 percent of low, medium, and high management potential, respectively (Table 4).

**Figure 3.**
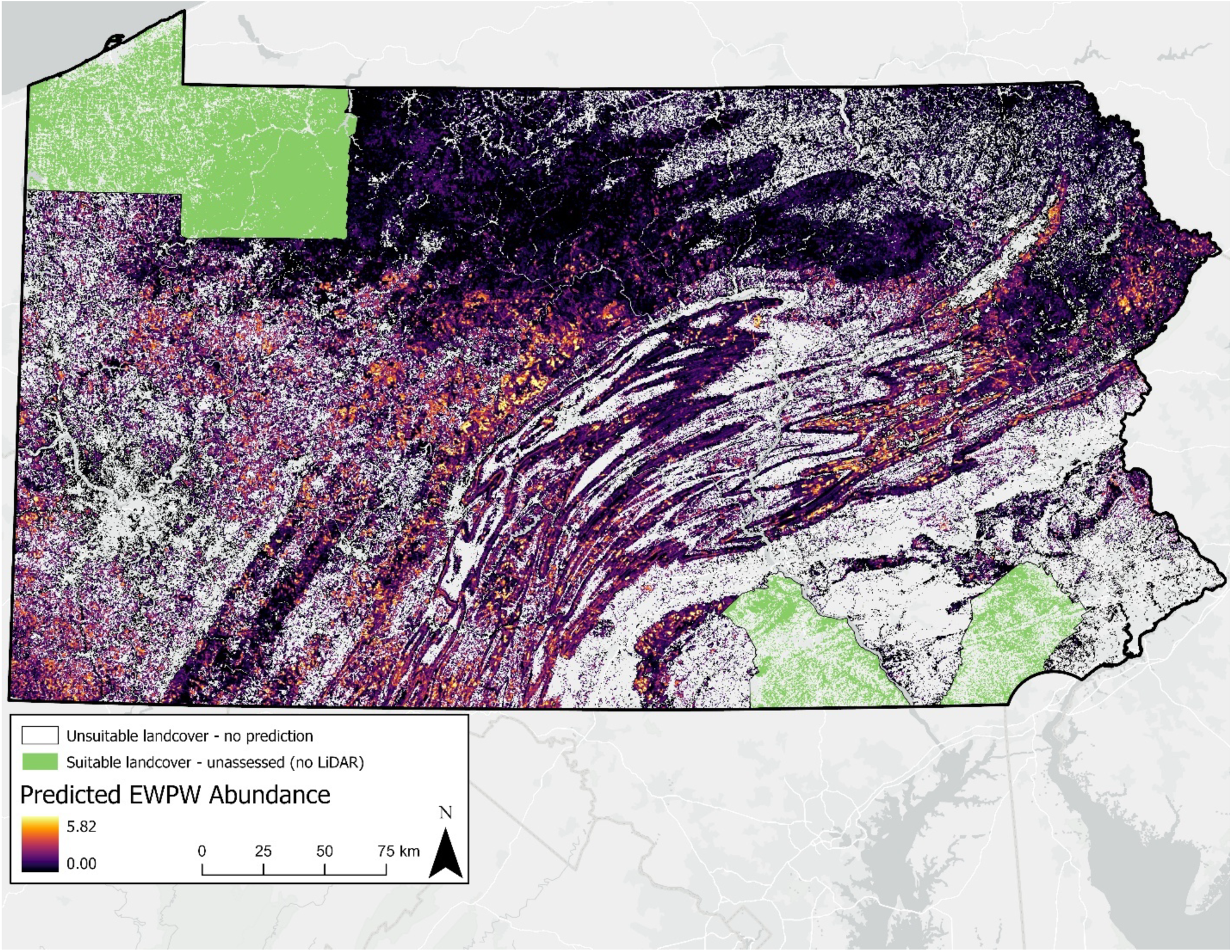
Map displaying a predicted eastern whip-poor-will (*Antrostomus vociferus*) abundance raster across Pennsylvania, USA in 2020 and 2021. Remotely sensed data (autonomous recording units, LiDAR, and landcover data) and a Royle-Nichols model was used to inform the raster. Abundance ranged from 5.82 (yellow) to 0.00 (black) individuals. White areas represent unsuitable land covers (e.g., agriculture or impervious). Green areas indicate where there was a suitable land cover (forest), but no LiDAR data to inform a prediction.

**Table 4.** Summary of forest area by ownership type based on two classified rasters: predicted abundance of eastern whip-poor-will (*Antrostomus vociferus*) and forest management potential. Each raster was classified into three categories: low, medium, and high. Columns 1–2 show forest ownership types and their total forested area. Column 3 presents the percentage of total forest area represented by each ownership type. Columns 4–6 report the area and percentage of forest within each ownership type classified as low, medium, and high. Columns 7–9 show the percentage of forest area within each ownership type falling into the three abundance classes.

| **Ownership Type** | **Forest cover (ha)** | **Forest ownership (%)** | **Low Abundance (ha; %)** | **Medium Abundance (ha; %)** | **High Abundance (ha; %)** | **Low Management Potential (%)** | **Medium Management Potential (%)** | **High Management Potential (%)** |
| --- | --- | --- | --- | --- | --- | --- | --- | --- |
| Private Land | 5,087,939 | 75.06 | 3512749; 73.14 | 1258500; 79.36 | 94726; 64.43 | 84.02 | 67.05 | 43.33 |
| State Forest | 897,856 | 13.25 | 700070; 14.58 | 163577; 10.32 | 26709; 18.17 | 7.23 | 19.69 | 32.17 |
| State Game Land | 567,199 | 8.37 | 398124; 8.29 | 142612; 8.99 | 23254; 15.82 | 5.49 | 10.06 | 20.50 |
| Federal Land | 131,045 | 1.93 | 113672; 2.37 | 8448; 0.53 | 1376; 0.94 | 2.44 | 1.18 | 0.82 |
| State Park | 94,779 | 1.40 | 78270; 1.63 | 12665; 0.80 | 950; 0.65 | 0.82 | 2.02 | 3.19 |

**Figure 5.**
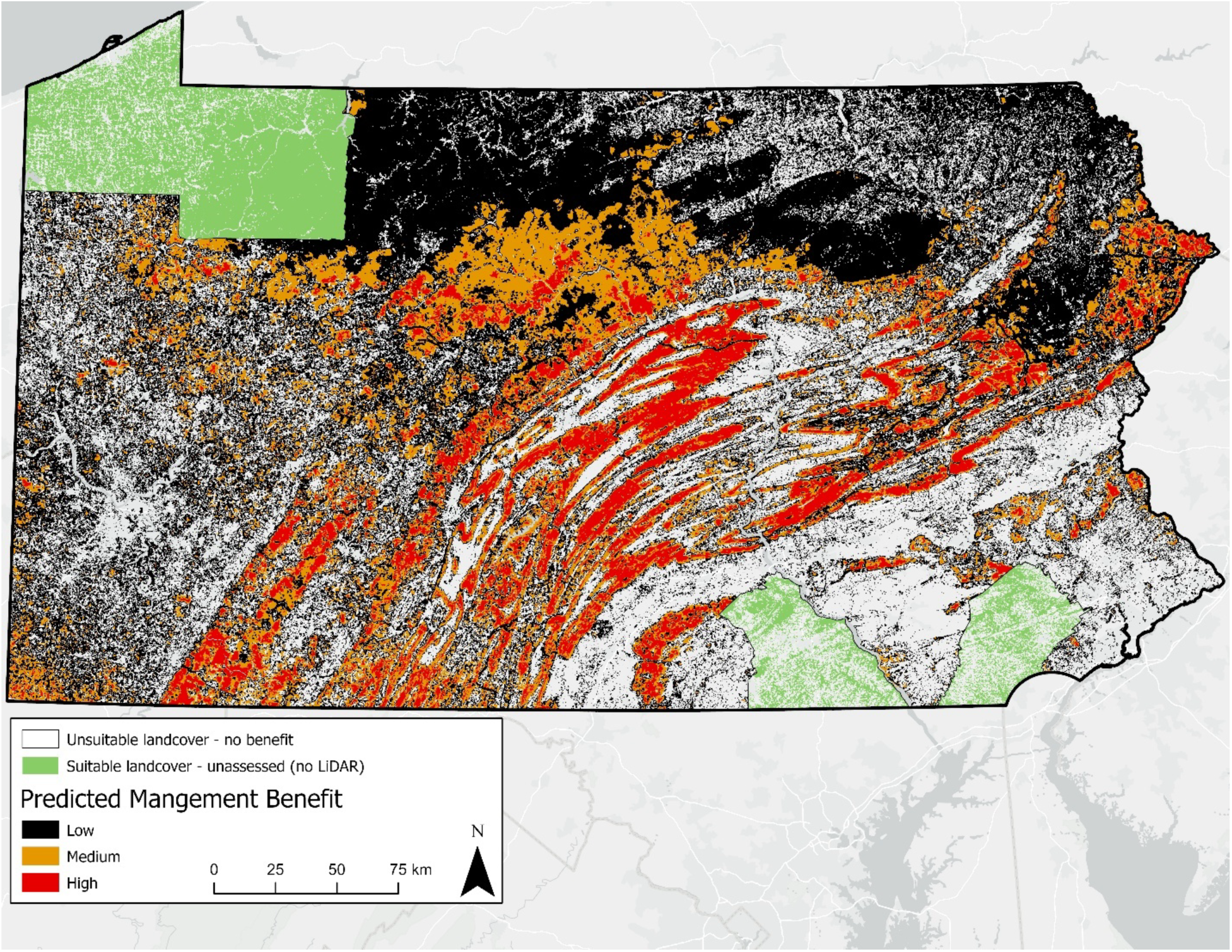
Map displaying a predicted eastern whip-poor-will (*Antrostomus vociferus*) management potential raster across Pennsylvania, USA in 2020 and 2021. Remotely sensed data (autonomous recording units, LiDAR, and landcover data) and a Royle-Nichols model were used to inform the raster. Management potential ranged from low (black), medium (orange), and high (red). White areas represent unsuitable land covers (e.g., agriculture or impervious). Green areas indicate where there was a suitable land cover (forest), but no LiDAR data to inform a prediction.

## Discussion

Our study provides valuable insight into landscape and forest structural conditions that influence whip-poor-will abundance. Further, we used model predictions to identify areas with high whip-poor-will abundance under current conditions and those where forest management that diversifies canopy structure has high potential to increase local abundance. Ultimately, these outcomes provide another example of how studies that incorporate data that quantify fine-scale forest structure (LiDAR) in addition to landscape composition (i.e., Dynamic World and USFS forest type) can better inform on-the-ground conservation for imperiled species (Farrell et al., 2013; Fricker et al., 2021). Such an approach can help land managers fine-tune stand-level management practices to best achieve desired structural conditions while also ensuring they are implemented within landscape contexts that are most attractive to the target species (Garabedian et al., 2017; Bombi et al., 2019; McNeil et al., 2023).

As hypothesized, whip-poor-will abundance was greatest at locations with high levels of horizontal complexity, which in part, stemmed from modest amounts of early successional forest. Both LiDAR-derived measures of forest structure included in our analyses, canopy height (p95) and the variation in canopy height (SD p95), influenced whip-poor-will abundance, which corroborates general conclusions from previous studies. For instance, whip-poor-will require early successional conditions, but are also known to use shelterwood establishment harvests and adjacent mature forests edges (Wilson and Watts, 2008; Thompson et al., 2022; Spiller et al., 2022). Estimated whip-poor-will abundance in our study peaked when forests within 300 m of a survey location had an average canopy height of 10.1 m and variation in canopy height of 7.4 m. To achieve these values simultaneously requires the presence of both early successional and mature forest within the local landscape. These conditions can be facilitated by ensuring regeneration harvests are proximate to mature forests, while also leaving scattered residual trees or retention islands within regeneration timber harvests. Previous studies have found whip-poor-will to be associated with such legacy features (Akresh et al., 2016; Wilson and Watts, 2008). Comparing LiDAR point clouds for survey locations with high and low whip-poor-will abundances provides managers with a visual perspective of the structural conditions they can emulate when implementing future habitat projects (Fig. 4).

While we identified structural elements that forest managers should consider when designing whip-poor-will habitat projects, our study also revealed compositional elements of the local landscape that were important drivers of whip-poor-will abundance. In support of our predictions, our analyses found whip-poor-will abundance was highest in heavily forested landscapes dominated by oak community types, with minimal impervious cover. The positive linear relationship we found between forest cover and whip-poor-will abundance is consistent with past research (Thompson et al., 2022; Souza-Cole et al., 2022). However, some of the most heavily forested portions of our study area (northern Pennsylvania) had the lowest predicted abundances, indicating that forest cover alone is not the sole landscape requisite for achieving high whip-poor-will abundance (Fig. 3). Indeed, in our study, forest type strongly influenced whip-poor-will abundance, whereby heavily forested landscapes dominated by oak community types host the highest predicted values. We postulate that this finding may be driven by differences in prey availability among forest types. It is well documented that Lepidoptera (i.e., moths) comprise the majority of whip-poor-will diets (Souza-Cole et al., 2022) and that oaks support more abundant and diverse Lepidopteran communities compared to other forest community types (Summerville and Crist, 2008; Narango et al., 2020). Thus, sites dominated by oak forest may host higher prey densities, which in turn could support higher whip-poor-will abundances. Collectively, our landscape context and forest structure findings stress the important interplay among forest area, community type, and canopy diversifying disturbances that create the conditions that support high whip-poor-will abundances. Such conditions may promote high abundances given individuals can meet their habitat needs (e.g., nesting, foraging, and roosting cover) within smaller home ranges compared to individuals inhabiting areas with less optimal conditions, a phenomenon observed in other bird species (i.e., Smith and Shugart, 1987; Bock and Jones, 2004; Diemer and Nocera, 2014).

We created rasters for predicted abundance and habitat management potential which can serve as valuable tools for informing whip-poor-will conservation (Figs. 3 and 5). For example, the likelihood of focal species occupying recently created habitat often depends on proximity to existing populations (Tittleret al., 2009; McNeil et al., 2020; Shaffer et al., 2025). As such, managers can reference the management potential raster to identify areas for future whip-poor-will habitat projects in landscapes with suitable conditions for the species, and then use the predicted abundance raster to prioritize projects based on their distance to areas that likely support existing populations. By using both rasters in combination, forest managers can more precisely implement management in locations that have the greatest potential benefit to whip-poor-wills. Comparatively, our predicted abundance raster aligns well with similar datasets for this species (e.g., Fink et al. 2023, Larkin et al. 2024b), but is more spatially explicit. For example, eBird (Fink et al. 2023) generates a whip-poor-will abundance map with estimates provided at a broad extent (2.5 km x 2.5 km resolution), whereas our map provides estimates at the sub-stand level (10 m x 10 m resolution). Ultimately, our rasters demonstrate that incorporating LiDAR and robust species occurrence data provides precision and resolution at scales that are ecologically meaningful to the target species and operationally meaningfully to land managers.

When reviewing these rasters in relation to forest ownership, conservationists can also identify stakeholders with forests that support existing whip-poor-will populations or forests that have strong potential to benefit the species via habitat management. Indeed, regional conservation efforts to recover declining species often depend on the collective efforts of multiple resource management agencies to meet desired outcomes (i.e., number of ha treated; Lott et al., 2021; White et al., 2023). Our analysis revealed that 64% of forests with high predicted abundances occurred on private land, which was to be expected given that private forests accounted for a significant percentage (75%) of forest cover included in our study. Nonetheless, this observation combined with the fact that private lands account for 43% of forests classified as having high management potential suggests that conservation programs that target private lands, like those offered by NRCS (Litvaitis et al., 2021), can play a major part in whip-poor-will recovery efforts.

While private forests clearly play an important role in present and future whip-poor-will conservation, the importance of public lands cannot be overstated. Public lands were estimated to support only 15% of eastern birds classified as forest obligates, but hosted >90% of two highly threatened species populations, the red-cockaded woodpecker (*Leuconotopicus borealis*) and Kirtland’s warbler (*Setophaga kirtlandii*; NABCI, 2011). Comparable to whip-poor-will, these two conservation dependent species require large forested areas with structural conditions maintained by periodic disturbances (Walters et al., 2002; Donner et al., 2008). Pennsylvania is fortunate to have more than 4 million acres of state-managed lands (PA DCNR, 2020), which are often interconnected and form expansive forested landscapes. Moreover, these lands have staff and financial resources to plan and implement forest management practices (e.g., timber harvest, forest stand improvements, and prescribed fire) that create and maintain a mosaic of stand age classes and structural conditions over time, which our analyses found to be important drivers of whip-poor-will abundance. As such, while forests managed by state agencies only accounted for 23% of the forest cover included in our analyses, they host a disproportionate amount of area classified as having high predicted abundance (35%) and high management potential (56%; Table 4). Moreover, these public lands contain the greatest concentration of areas with the highest predicted abundances, which represent strong population anchors for regional habitat-based conservation efforts. Ultimately, examining our predicted abundance and management potential rasters through the lens of forest ownership suggests that multi-agency management actions on both public and private land have the potential to contribute meaningfully toward achieving statewide conservation goals for this imperiled species. Given that our models clearly illustrate the importance of transient forest conditions (i.e., low canopy height), conservation planning across these broad ownerships will require spatial and temporal considerations to ensure these conditions are being created across the landscape to support the species long-term.

## Conclusions

When considered together, our results indicate that whip-poor-will abundance is highest in heavily forested, oak-dominated landscapes with complex age class and structural conditions created through natural processes (e.g., forest succession) and periodic disturbances (both natural or anthropogenic; e.g., tornados, timber harvest, and prescribed fire). Current eastern forests are experiencing unprecedented challenges that are leading to simplification, degradation, and mesophication (e.g., invasive species, intense ungulate browsing, high-grading, and suppression of disturbances; Knoot et al., 2010; Dey, 2014). Without intervention, the compositional and structural conditions that whip-poor-will require will remain scarce in eastern forests. Managers who seek to create habitat for whip-poor-will through a natural communities lens should consider restoration and maintenance of woodland systems (e.g., open oak woodlands; Dey et al., 2017), a forest community dominated by oak species that once covered more than 100 million ha of eastern North America (Hanberry et al., 2020). Woodland systems, like any forest, experienced forest dynamics (e.g., heterogeneous fire return intervals and forest succession) which resulted in variable structure (Dey et al., 2017; Hanberry et al., 2018). Our results suggest that managers who seek to balance timber production, while also benefiting wildlife, may consider implementing a shelterwood system that targets oak with various stages of the shelterwood sequence (preparatory, establishment, and removal harvests) interspersed across a landscape at any given time (Loftis, 1990; Ashton and Kelty, 2018). Indeed, the restoration/maintenance of woodlands or the implementation of shelterwood systems would be most impactful in landscapes we identified as having high management potential (Fig. 5).

The information presented here, in addition to past research, provides managers with a well-grounded understanding of the conditions needed to support breeding whip-poor-will. We recommend future conservation efforts for whip-poor-will involve leveraging publicly available LiDAR datasets to identify actionable management targets and create tools that facilitate habitat management, such as those produced by our study (Figs. 3 and 5). There remains much insight to be gained from studies that incorporate fine-scale forest structure data to examine other aspects of whip-poor-will ecology and demography. For example, studies that use LiDAR-derived covariates to explore how nesting and post-fledging survival is influenced by fine-scale forest structure would be valuable. Such studies may provide important information regarding the structural conditions associated with high quality nesting and post-fledging habitat, in addition to their optimal spatial arrangement, which can be incorporated into future conservation efforts for this species. Lastly, we encourage the conservation community to continue to incorporate LiDAR-derived environmental variables into studies that aim to elucidate habitat relationships or to inform spatially explicit conservation plans for other species or groups of species. As demonstrated here, LiDAR data can substantially contribute to understanding and mapping species distributions and other demographic parameters at high resolutions. If recent LiDAR datasets are unavailable, collaborations to fund data acquisition would be worthwhile.

## Acknowledgements

We would like to thank the many private landowners and field technicians that made this work possible. We are also grateful to the Pennsylvania Game Commission and Pennsylvania Department of Conservation and Natural Resource-Bureau of State Parks for land access and logistics. We are grateful for the support of the bridges2 (psc.edu/resources/bridges-2/user-guide) and Unity (https://unity.rc.umass.edu/) supercomputing clusters. Finally, we are thankful for the insight and suggestions provided by the two anonymous reviewers. The views and conclusions contained in this document are those of the authors and should not be interpreted as representing the opinions or policies of the U.S. Government or the National Fish and Wildlife Foundation and its funding sources.

## CRediT authorship contribution statement

Jeffery T. Larkin: Conceptualization, Methodology, Investigation, Formal analysis, Data curation, Writing – original draft, Writing – review & editing; Malcolm Itter: Funding, Formal analysis, Writing – original draft, Writing – review & editing; Cameron J. Fiss: Investigation, Formal analysis, Writing – review & editing; Lauren M. Chronister: Software, Data curation, Writing – review & editing; Justin Kitzes: Data curation, Writing – review & editing; Jeffery L. Larkin: Funding acquisition, Project administration, Conceptualization, Methodology, Writing – review & editing; Halie Parker Larkin: Investigation, Writing – review & editing; Darin J. McNeil: Formal analysis, Writing – original draft, Writing – review & editing; Anthony W. D’Amato: Writing – review & editing; Michael E. Akresh: Funding acquisition, Writing – review & editing; David I. King: Funding acquisition, Project administration, Writing – review & editing

